# Classification of complex emotions using EEG and virtual environment: proof of concept and therapeutic implication

**DOI:** 10.1101/2020.07.27.223370

**Authors:** Eleonora De Filippi, Mara Wolter, Bruno Melo, Carlos J. Tierra-Criollo, Tiago Bortolini, Gustavo Deco, Jorge Moll

## Abstract

During the last decades, neurofeedback training for emotional self-regulation has received significant attention from both the scientific and clinical communities. However, most studies have focused on broader emotional states such as “negative vs. positive”, primarily due to our poor understanding of the functional anatomy of more complex emotions at the electrophysiological level. Our proof-of-concept study aims at investigating the feasibility of classifying two complex emotions that have been implicated in mental health, namely tenderness and anguish, using features extracted from the electroencephalogram (EEG) signal in healthy participants. Electrophysiological data were recorded from fourteen participants during a block-designed experiment consisting of emotional self-induction trials combined with a multimodal virtual scenario. For the within-subject classification, the linear Support Vector Machine was trained with two sets of samples: random cross-validation of the sliding windows of all trials; and 2) strategic cross-validation, assigning all the windows of one trial to the same fold. Spectral features, together with the frontal-alpha asymmetry, were extracted using Complex Morlet Wavelet analysis. Classification results with these features showed an accuracy of 79.3% on average when doing random cross-validation, and 73.3% when applying strategic cross-validation. We extracted a second set of features from the amplitude time-series correlation analysis, which significantly enhanced random cross-validation accuracy while showing similar performance to spectral features when doing strategic cross-validation. These results suggest that complex emotions show distinct electrophysiological correlates, which paves the way for future EEG-based, real-time neurofeedback training of complex emotional states.

**Significance statement:** There is still little understanding about the correlates of high-order emotions (i.e., anguish and tenderness) in the physiological signals recorded with the EEG. Most studies have investigated emotions using functional magnetic resonance imaging (fMRI), including the real-time application in neurofeedback training. However, concerning the therapeutic application, EEG is a more suitable tool with regards to costs and practicability. Therefore, our proof-of-concept study aims at establishing a method for classifying complex emotions that can be later used for EEG-based neurofeedback on emotion regulation. We recorded EEG signals during a multimodal, near-immersive emotion-elicitation experiment. Results demonstrate that intraindividual classification of discrete emotions with features extracted from the EEG is feasible and may be implemented in real-time to enable neurofeedback.

## Introduction

“This world’s anguish is no different from the love we insist on holding back,” wrote Aberjhani in the book “Elemental: The Power of Illuminated Love” published in 2008 with the prize-winner artist Luther E. Vann. A decade later, there is still much to discover about the neurophysiology of complex emotions, which, according to some, represent a combination of basic emotions (Barrett et al., 2007) or primary emotive states and cognitive components such as event-feature-emotion complexes (Moll et al., 2005; 2008). The purpose of our work is to explore to which extent electrophysiological activity related to one specific affiliative emotion (Moll et al., 2012; Cacioppo & Cacioppo, 2012), tenderness/affection, can be distinguished from anguish. We focus on tenderness as an affiliative feeling because it is the basis for empathy and prosocial behavior (Eslinger, 1998; Zahn et al., 2009), being associated with social bonding, care, and wellbeing. Anguish, to the contrary, reflects a negative state of mental suffering linked to social dysfunction, withdrawal, and poor mental health (Corbett, 2015).

Given the importance of affiliative emotions for a range of psychosocial processes (Baumeister & Leary, 1995), psychotherapeutic approaches have embraced the training of such emotions as part of the therapeutic process in the form of compassion-focused therapy (Gilbert, 2006; Germer et al., 2012) and loving-kindness meditation (Lutz et al.,2008; Neff et al., 2013).

Real-time fMRI-Neurofeedback (NFB) training has also been broadly explored concerning the self-regulation of emotions. While most of NFB experiments have been focusing on regulating the activity of the amygdala (Zotev et al., 2011, 2018; Young et al.,2014, 2017; Paret et al., 2014), Lorenzetti et al. (2018) demonstrated that the self-regulation of anguish and tenderness is also possible by targeting the septohypothalamic network, previously found to be engaged in affiliative feelings (Moll, 2011;2012; Zahn, 2009b). Although fMRI has the advantage of a higher spatial resolution and allows modulating deep brain structures, it also has its downsides, such as the signal delay (Friston, 1994), an environment that can be hostile and uncomfortable, and high costs of scanning. In contrast, EEG offers a higher temporal resolution and a direct measure of information processing. Moreover, it is cheaper and rather simple in use, which makes it a more accessible tool in the clinical environment. Research in emotion classification with EEG has mostly focused on distinguishing emotions based on psychological models (Russell, 2003; Rubin et Talarico, 2009), primarily using the “circumplex model” (Lang et al., 1993), which portrays different types of emotions in a two-dimensional space comprising valence and arousal. Another model, the discrete emotional model, has mainly been explored with regards to basic emotions (i.e., sadness, happiness, fear, surprise, anger, and disgust) (Davidson, 1992; Balconi and Lucchiari, 2006; Baumgartner et al., 2006; Balconi and Mazza, 2009; Li and Lu, 2009).

Therefore, our study aims at establishing a method for classifying complex emotions that can be later used for EEG-based neurofeedback on emotion regulation.

Attempts to differentiate tenderness from negative emotions using EEG have been made, as in the two studies conducted by Zhao et al. (2018;2018). In these studies, the authors found the frontal alpha asymmetry (FAA) and the midline theta power to be useful features to classify such emotional states. Therefore, we included these features in our analysis, combined with other spectral and correlational measures across the whole scalp. We hypothesized that tenderness and anguish have different electrophysiological correlates, not reduceable just to the FAA, that can be distinguished using a machine-learning approach.

A significant difference in our study lies in the way we evoked emotions. We employed a multisensory virtual environment together with musical excerpts, in contrast to single-modality stimuli (e.g., videoclips) used in previous studies (Zhao et al., 2018; 2018). Moreover, we encouraged participants to use personalized “mantras” (i.e., short emotionally loaded sentences) to facilitate self-induced emotional states. This approach to emotion induction can, indeed, be of great advantage when using neurofeedback as a self-regulation training (Yankauer, 1999).

## Materials and methods

### Participants

For this proof of concept experiment, 14 healthy adults or young adults (7 males and 7 females) with no history of psychiatric disorder or neurologic illness were recruited through advertisements both on the internet and at universities.

EEG recordings from three participants (2 males and 1 female) were discarded before analysis because of excessive artifacts due to equipment malfunctions. The average age of the 11 remaining participants was 27 years (SD = 7), ranging from 19 to 46. All participants had a normal or corrected-to-normal vision and had Brazilian nationality, except one who was German but fluently spoke Portuguese.

### Emotion-inducing stimuli

We implemented a multimodal stimulation with a virtual scenario, an edited version of the Unity 3D asset Autumnal Nature Pack, accompanied by emotion-specific musical excerpts. The virtual environment displayed the first-person view over a landscape of hills and cornfields, differently colored according to the emotion elicited: purple for anguish and orange for tenderness (Fig.1). Together with these visual stimuli, participants were simultaneously listening to eight (four for each of the two emotions) different instrumental musical excerpts of 46-seconds. Music excerpts were fixed for each trial type (i.e., 4 for anguish and 4 for tenderness) and normalized with the root mean square feature on the software Audacity (Audacity, http://www.audacityteam.org). For all tenderness trials, a piece of mild, harmonic, and gentle music was used. In contrast, the unpleasant stimuli for the anguish trials were electronically manipulated from the original pleasant tunes used for the tenderness condition. For each pleasant stimulus, a new sound-file was created, in which the original (pleasant) excerpt was recorded simultaneously with two pitch-shifted versions of the same excerpt. The pitch-shifted versions were one tone above and a tritone below the original pitch (samples of the stimuli are provided at http://www.stefan-koelsch.de/Musiction1), resulting in dissonant and distorted music. For the neutral condition, the landscape was typically colored, and no background music was presented.

### Experimental procedure

Upon arrival, all participants signed the informed consent and filled up the State-Trait Anxiety Inventory (*STAI*), the Beck’s Depression Inventory (BDI), and The Positive and Negative Affect Schedule (*PANAS*). After the EEG cap setup, participants were instructed to use mantras and personally recalled memories to facilitate emotional self-induction during the experiment. As guidance, we provided a list of suggested mantras (Table 1), but they were free to choose mantras personally. Throughout the experiment, subjects were comfortably seated in an armed chair approximately 50 cm away from the screen, wearing padded headphones. The experimental procedure consisted of 8 emotion-alternating blocks (4 for anguish trials and 4 for tenderness), as represented in Figure 2. Each block included four emotion-eliciting trials (46 s) interleaved by four short neutral trials (12 s), which allowed for a brief rest between each emotional induction. The auditory and visual stimuli used for each trial are the ones described in the previous section. At the end of each block, participants had to fill a short questionnaire and rate the emotional intensity they felt, the concentration, and fatigue levels. We recorded EEG signals throughout the whole experiment, and no break was given between the blocks.

**Table 1.** Samples of mantras used according to the experimental conditions. Free translation from Portuguese to English is provided below each sentence in italics.

| **Tenderness** | **Anguish** | **Neutral** |
| --- | --- | --- |
| Compaixão protege tudo | Não vou conseguir nada | A Terra é redonda |
| *Compassion protects everything* | *I will not achieve anything* | *The earth is round* |
| Sentir o cheiro de um bebê<br>*The smell of a baby* | Doenças afligem tudo<br>*Diseases afflict everything* | A grama está crescendo<br>*The grass is growing* |
| Harmonia está em toda parte<br>*Harmony is everywhere* | Violência está em toda parte<br>*Violence is everywhere* | A mesa é marrom<br>*The table is brown* |
| Amor está em toda parte<br>*Love is everywhere* | O mundo está cheio de ódio<br>*The world is full of hate* | A luz está acesa<br>*The light is on* |

**Figure 1:**
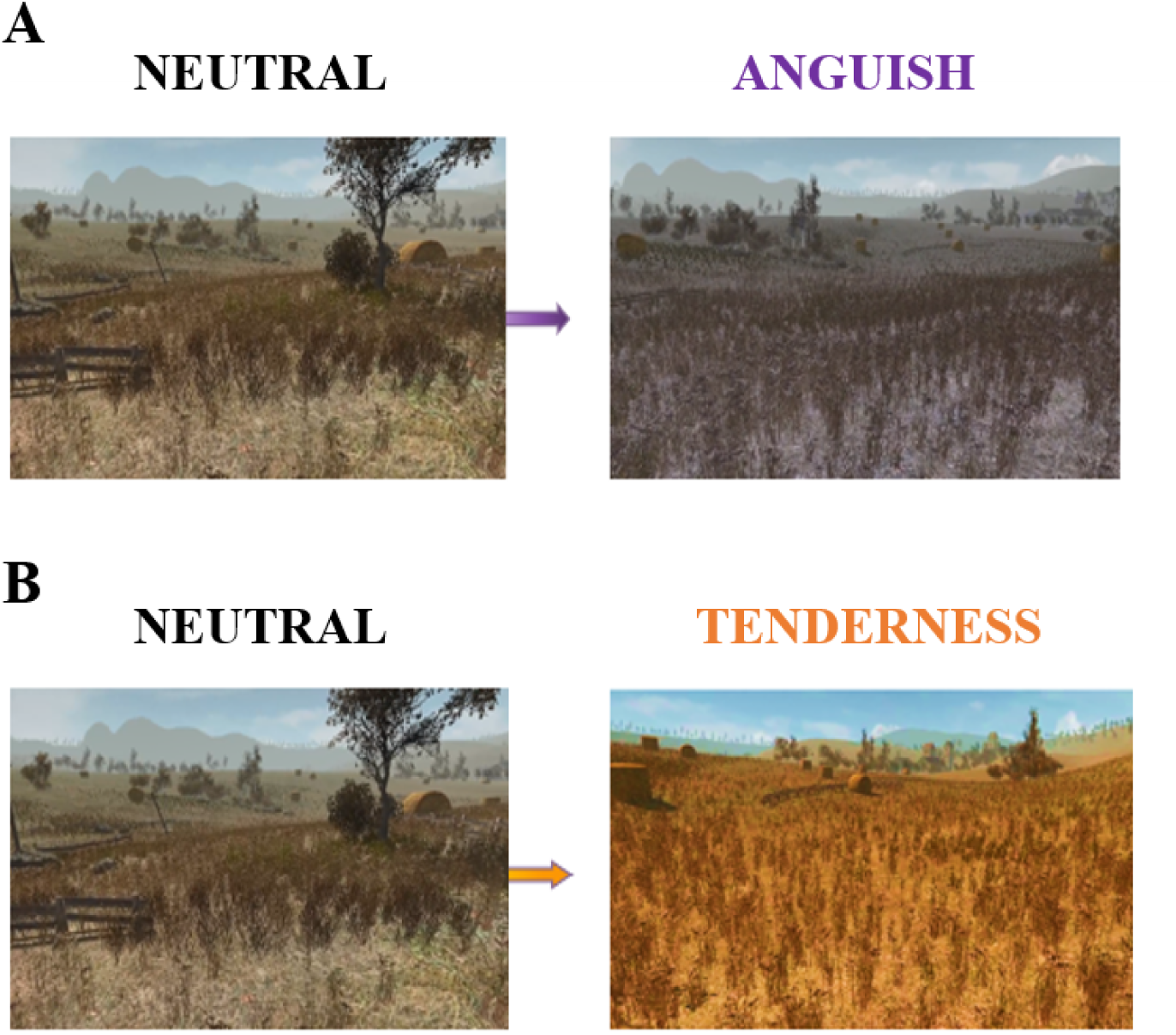
The virtual reality scenario used to induce emotions during the experiment. Compared to the neutral colored scene on the left, the color hue of the landscape turned into purple (Fig.1A) for anguish trials, while for all tenderness trials, it turned into a vivid orange (Fig. 1B).

**Figure 2:**
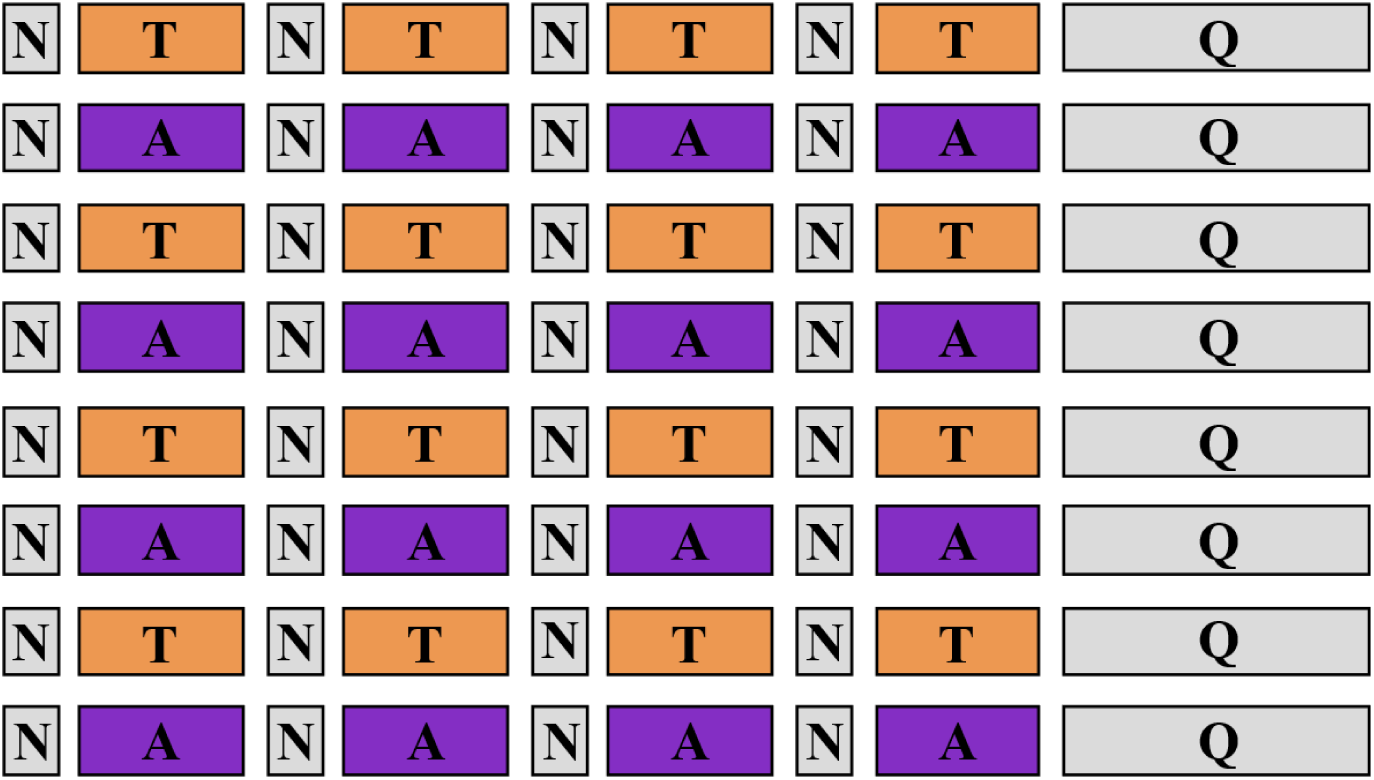
Experimental design. Eight emotion-alternating blocks (orange boxes refer to “tenderness” condition, and purple boxes to “anguish”), each consisting of 4 emotion-inducing trials of 46s, interleaved by 4 short neutral trials of 12s. At the end of each block, a self-assessment questionnaire was filled out by the participants.

### EEG acquisition

EEG data were recorded using the Brain Vision Recorder software, and the signal was acquired from the BrainAmp amplifier and the standard 64 channels EEG-MR Plus cap (Brainproducts, Germany) at a sampling rate of 1000 Hz and a bandwidth of 0,01-250 Hz. All the electrodes had sintered Ag/AgCl sensors placed according to the standardized international 10/20 EEG placement system, and their impedance was kept under seven kl. One out of the 64 channels was used to record the ECG signal, while the reference electrode was channel FCz and the ground channel AFz. The experiment pipeline was programmed on Matlab (The Mathworks, Inc.).

### EEG preprocessing

Offline analysis of EEG data was done using the EEGlab toolbox (Delorme et al.,2004) on Matlab R2019b (The Mathworks, Inc.). The preprocessing pipeline firstly included downsampling the signal to 250 Hz and applying a bandpass Butterworth filter of 0.01-45 Hz. The Independent Component Analysis (ICA) algorithm was applied to correct for eyeblinks and muscular artifacts. Components which captured artifacts were manually pruned, independently for each subject. After artifact removal, the EEG dataset was epoched and cut into three distinct datasets for each subject according to the experimental condition. For each participant, we created one dataset for the 16 tenderness trials of 46s length each, one for all the 16 anguish trials of 46s length each, and one containing all 32 neutral trials of 12s length. Data were then visually inspected, but no further artifact or epoch rejection method was applied. This choice was based on our future goal of using a similar setup for real-time neurofeedback studies, which will preclude the possibility to visually inspect and reject bad epochs. Therefore, we pursued a method allowing for effective classification despite the presence of some residual artifacts.

Finally, we applied the surface Laplacian transform by implementing algorithms (Cohen, 2014) inspired by the spherical spline method described by Perrin et al. (1987a, 1987b, 1989). This spatial filter allows reducing volume conduction effects for connectivity analysis purposes.

### Feature extraction

#### Time-frequency analysis

EEG data were transformed into the time-frequency domain by using Complex Morlet Wavelet convolution in order not to lose information about the temporal dynamics.

The Complex Morlet wavelet (CMW) is a complex-valued sine wave tapered by a Gaussian window described by the following equation:

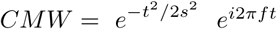

Where 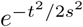 is the real-valued Gaussian and *e*^*i*2*πft*^ is the result of the Euler’s formula combined with a sine wave (Cohen, 2019). The time *t* is centered with regards to the wavelet by taking -2: sampling rate:+2 to avoid phase shifts.

One of the benefits of the CMW convolution over other methods such as Short-Fast Fourier Transform or the Hilbert Transform is the Gaussian-shaped wavelet in the frequency-domain. However, the caveat when doing convolution with a Complex Morlet Wavelet is to accurately define the width of the Gaussian, here defined as *s*, a key parameter for determining the trade-off between temporal and spatial resolution of the time-frequency analysis (see Mike X Cohen, 2019).

The parameter *s* is expressed as *s* = *c /*2*πf*, where*c* denotes the number of cycles of the wavelet, which is dependent on the frequency *f* of the same. A narrower Gaussian with fewer cycles in the time domain leads to a high temporal resolution but reduced spectral precision, and vice-versa with a wider Gaussian. Therefore, we applied a variable number of cycles ranging from 3 up to 10, increasing as a function of frequency to have a balanced trade-off between temporal and spectral resolutions. Since we were interested in all frequency bands, we selected a range of frequencies going from 1 up to 45 Hz. After applying CMW convolution, we extracted the power from the coefficients, and we applied a decibel-baseline normalization. Considering that we were mainly interested in changes in spectral features from a neutral mental state to the two distinct emotional ones, we used all the neutral trials as a baseline. To increase the number of samples, we cut the time-frequency data of each trial with a sliding-windows approach. Each window was 1 second long with a half-second overlap, resulting in a total of 91 windows per trial.

Then, the average change in power compared to the neutral baseline was calculated for seven frequency bands (delta 1-4 Hz, theta 4-8 Hz, low alpha 8-10 Hz, high alpha 10-12 Hz, low beta 13-18 Hz, high beta 18-30 Hz, and gamma 31-45 Hz). Within each window and for all the 63 channels and the seven frequency bands, the features extracted were the mean power, the standard deviation of the mean, and the frontal-alpha asymmetry (FAA). The FAA coefficients were calculated for the channel pairs Fp1-Fp2 and F3-F4 in both low-alpha (8-10 Hz) and high-alpha (10-12 Hz) bands. The resulting feature array consisted of 1456 windows as samples for each of the two classes (anguish and tenderness) with a total of 886 features.

#### Amplitude time-series correlation

After CMW, we extracted amplitude information of each channel and for each frequency component in the range 1-45 Hz.

We then applied the sliding-window approach again, so within each of the 91 windows and for each of the seven frequency bands aforementioned, we calculated the Spearman correlation coefficient of the 63*63 channels matrix. This time-frequency correlational analysis allows characterizing both the time-varying and frequency-varying properties of non-stationary signals such as electrophysiological signals. To eliminate redundant information and reduce the size of the feature array, we only selected all the coefficient in the upper triangle of the correlational matrices. The resulting high-dimensional feature array consisted of 13672 features (pairwise channels correlation coefficient for each frequency band) and again 1456 windows as samples for each of the two emotions.

#### Classification methods

For classification and visualization of the data, we used the MATLAB R2019b Statistics and Machine Learning Toolbox. We opted for a linear support-vector-machine (SVM) algorithm for high-dimensional data for binary classification of the feature arrays. SVMs are supervised learning algorithms that define a hyperplane as a decision boundary, such that the margin of separation between two classes is maximized. Herewith, SVMs provide a measure that allows scaling the certainty to which a window sample is assigned to one of the two classes: the distance of the sample from the separating hyperplane. Regarding future applications in NFB, this allows for a scaling of the feedback (e.g., gradually changing color hue) and, thus, more precise response to changes in emotional intensity.

We used two methods to train and validate the classifier. Firstly, we performed 10-fold cross-validation across all the window samples. In*k* -fold cross-validation, the samples are randomly partitioned into *k* equally sized sets. The classifier is trained *k* times, each time holding a different set out for validation. As a result, the algorithm returns *k* separately trained models. The classification accuracy for each of the models is determined by classifying the held-out fold. The accuracy returned for the classifier is the average of these results. We iterated 10-fold cross-validation ten times and subsequently averaged across classification results. In the following, we will refer to this validation method as random cross-validation.

Secondly, we applied 8-fold cross-validation in consideration of the trials, whereby all windows of one trial were assigned to the same fold. The windows of four trials (2 ‘Tenderness’, 2 ‘Anguish’), were thus kept as validation sets, while the classifier was trained with the remaining 28 trials. Again, we iterated the 8-fold cross-validation ten times and averaged across classification runs. Thinking about possible future applications in NFB studies, we chose this method to estimate the impact of event-specific markers on classification accuracy. As the model will be trained prior to the NFB session, we aimed at testing its performance on an entirely unknown set of data. Throughout the paper, we will refer to this method as strategic cross-validation.

To visualize the datasets, we used t-Distributed Statistic Neighbour Embedding (t-SNE) (Van Der Maaten & Hinton, 2008). This technique plots the high-dimensional data into a low-dimensional space by maintaining the basic structure of the dataset, such as clustering or blending of the samples from the two emotional states.

Feature-Ranking was performed with the Bioinformatics toolbox of MATLAB 2019b. Using the t-test as an independent criterion for binary classification, we ranked the features after their significance between the classes. The built-in function determines the absolute value of the two-sample t-test with pooled variance estimate for each feature. Essentially, the algorithm estimates how unlikely it is for a given feature, that the difference in the mean and variance of both emotional states occurred by chance. We extracted the 20 highest ranked features for each participant and feature array and subsequently evaluated results across participants.

## Results

### Subjective ratings

The intensity of the emotions, usefulness of mantras, fatigue, and concentration levels were assessed at the end of each block. An overview of subjective ratings is presented in Figure 3. Participants had to rate the intensity of the emotions on a five-point Likert scale going from “very mild” up to “highly intense.” On average, the emotional intensity was rated “moderate” to “high” for the first and the last blocks, with a slight increase for the second and the fourth blocks. As shown in Figure 3, there is no difference between the two emotions, suggesting that participants felt both tenderness and anguish with the same intensity. Concentration and fatigue levels were also self-rated on a five-point Likert scale going from “very low” up to “very high.” Concentration levels show minor differences across conditions and blocks, with the highest concentration level reported during the first block for both emotions. As expected, fatigue ratings showed a slightly increasing trend throughout the experiment, with a “very low” and “low” level for the first blocks and up to “moderate” levels for the last blocks. The usefulness of mantras was also rated on a 5-point Likert scale going from “not useful” up to “very useful”. Averaged responses across participants showed higher variance, compared to the other self-rated measures, across blocks and conditions, as shown in Figure 3D.

**Figure 3:**
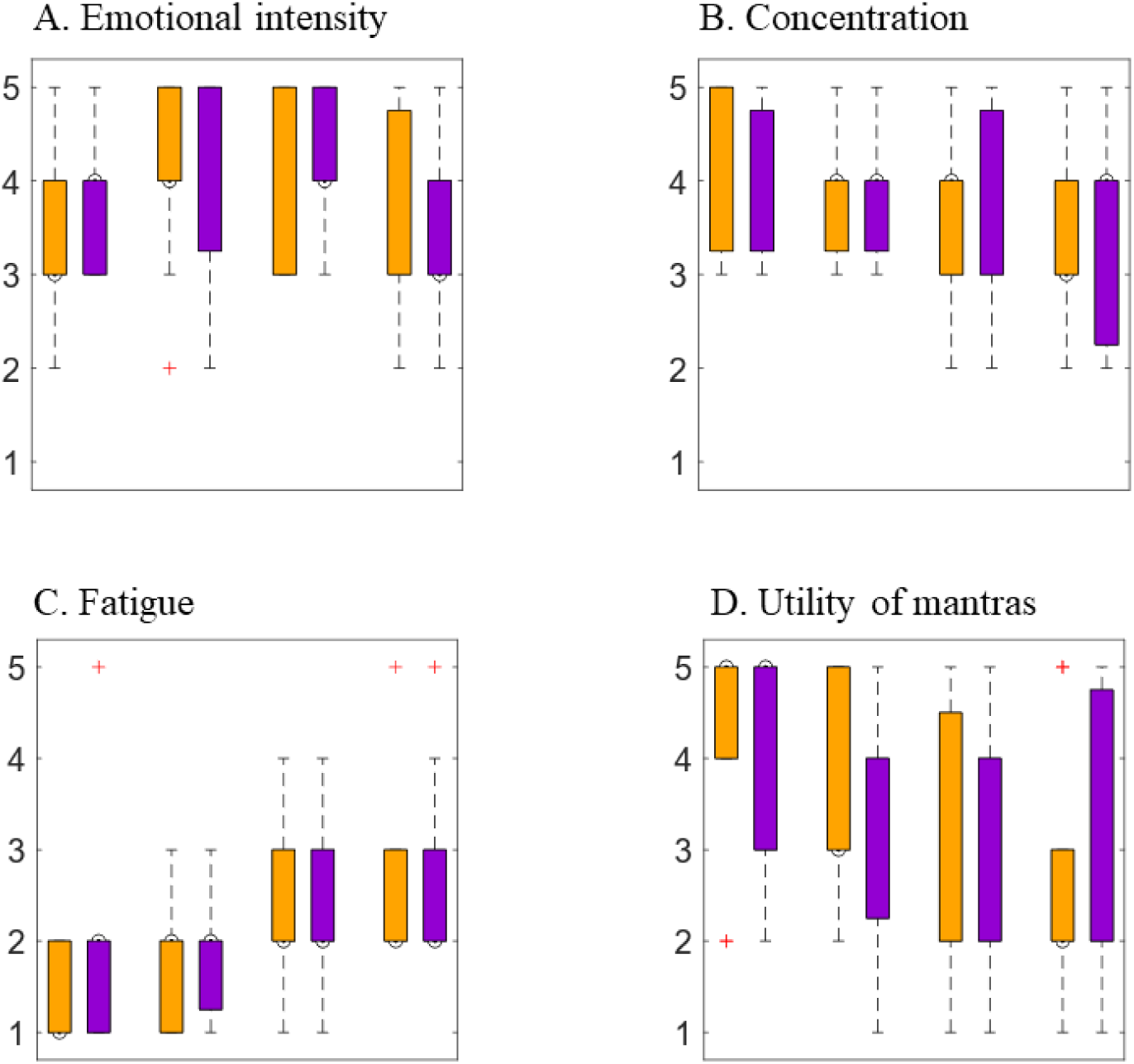
Participants’ subjective ratings assessed at the end of each of the eight blocks (four for each emotion). Orange refers to the “tenderness” condition, while purple refers to “anguish”. The y-axis represents the 5-point Likert scale, going from “very mild” to “very intense” for A, from “very low” up to “very high” for B and C and from “not useful” to “very useful” for D.

### Classification results using SVM

Ten independently cross-validated classification results for each participant with the described feature extraction and validation methods are presented in Figure 4. Corresponding average accuracies are summarized in Table 2. For the features extracted with the time-frequency analysis and FAA, we yield accuracies ranging from 66.3% up to 95.6% for the random validation sets (Fig. 4A) and 59.2% up to 92.9% when applying the strategic cross-validation method (Fig. 4B). The accuracy drops by 6.0% on average, when the validation samples originate from trials entirely unknown to the classifier. For the feature set extracted with amplitude time correlation analysis, we can report accuracies of 91.9 - 99.2 % for random cross-validation (Fig. 4C) and 62.4 - 92.4% for strategic cross-validation (Fig. 4D), which corresponds to an average 22.3% drop in the accuracy. These results show a high inter-subject variability across all feature sets and cross-validation methods.

**Table 2.**
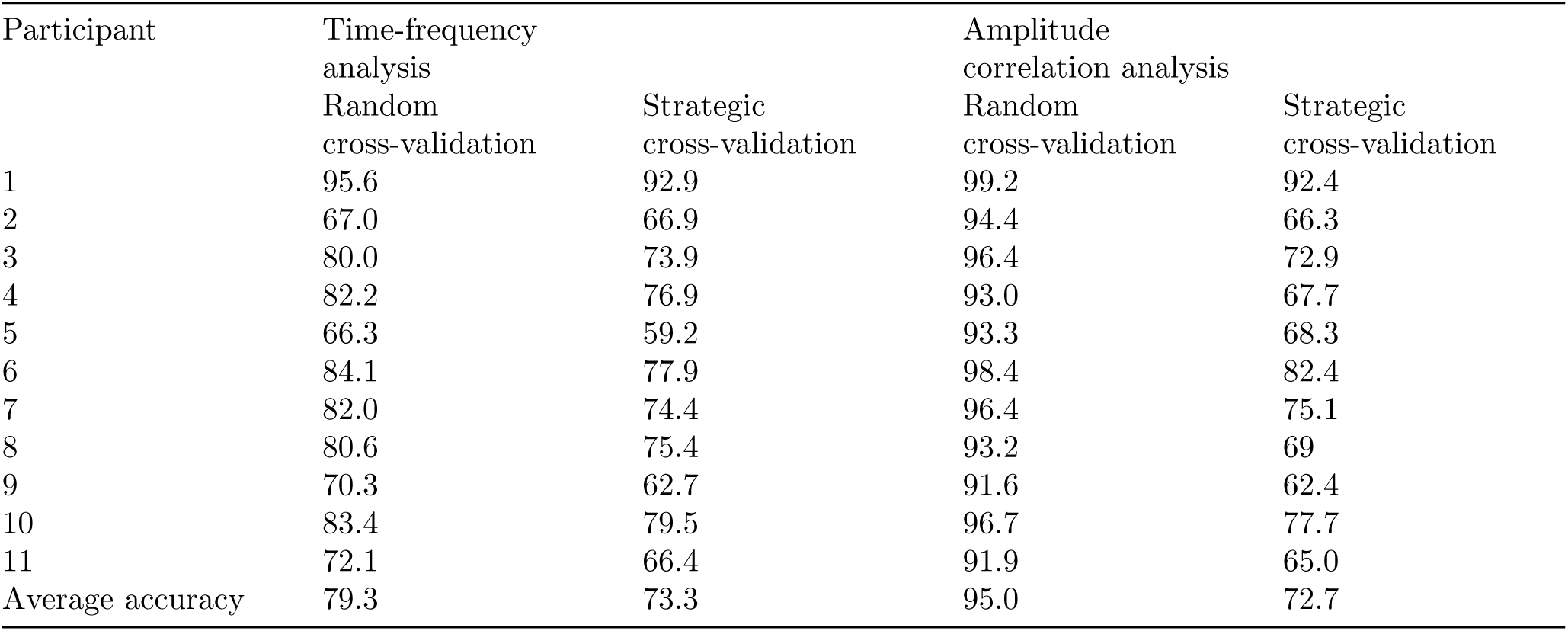
Classification results in percent for each participant, feature extraction, and cross-validation method.

**Figure 4:**
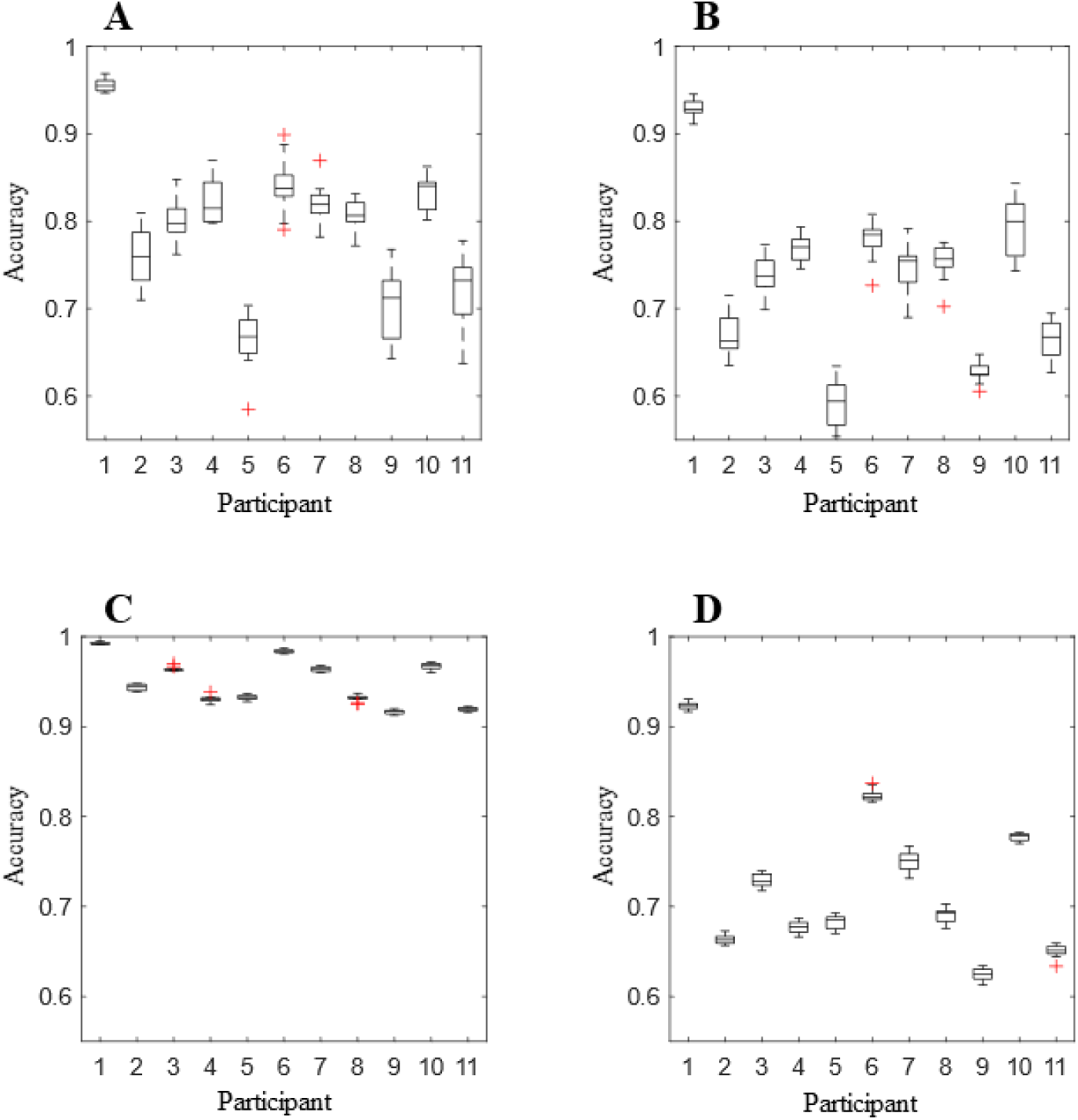
Classification results for each feature extraction and cross-validation methods. In the two upper graphs (A and B), classification was done with the features extracted from the time-frequency and FAA analysis, while the lower two graphs (C and D) display the results with the amplitude time-series correlation features. In boxplots A and C, we randomized all window-samples and performed 10-fold cross-validation. In boxplots B and D, we performed 8-fold cross-validation, whereby each validation fold consisted of the windows of 4 trials that were entirely unknown to the classifier.

To understand the variability of classification results, we plotted the datasets from the participants with the best and worst classification accuracy with t-SNE. The global geometry of the datasets with the best accuracy (Fig. 5A and C) can be separated into two clusters corresponding to the emotional states, while the same global structure cannot be found for the datasets with the worst accuracy (Fig. 5B and D). Notably, the datasets resulting from the amplitude time correlation analysis (Fig. 5C and D), display a more defined regional pattern, namely a clustering of samples into lines. These clusters consist of initially temporally adjacent samples. For the participant with the worst classification accuracy, this regional pattern dominates over the global data structure (Fig. 5D).

**Figure 5:**
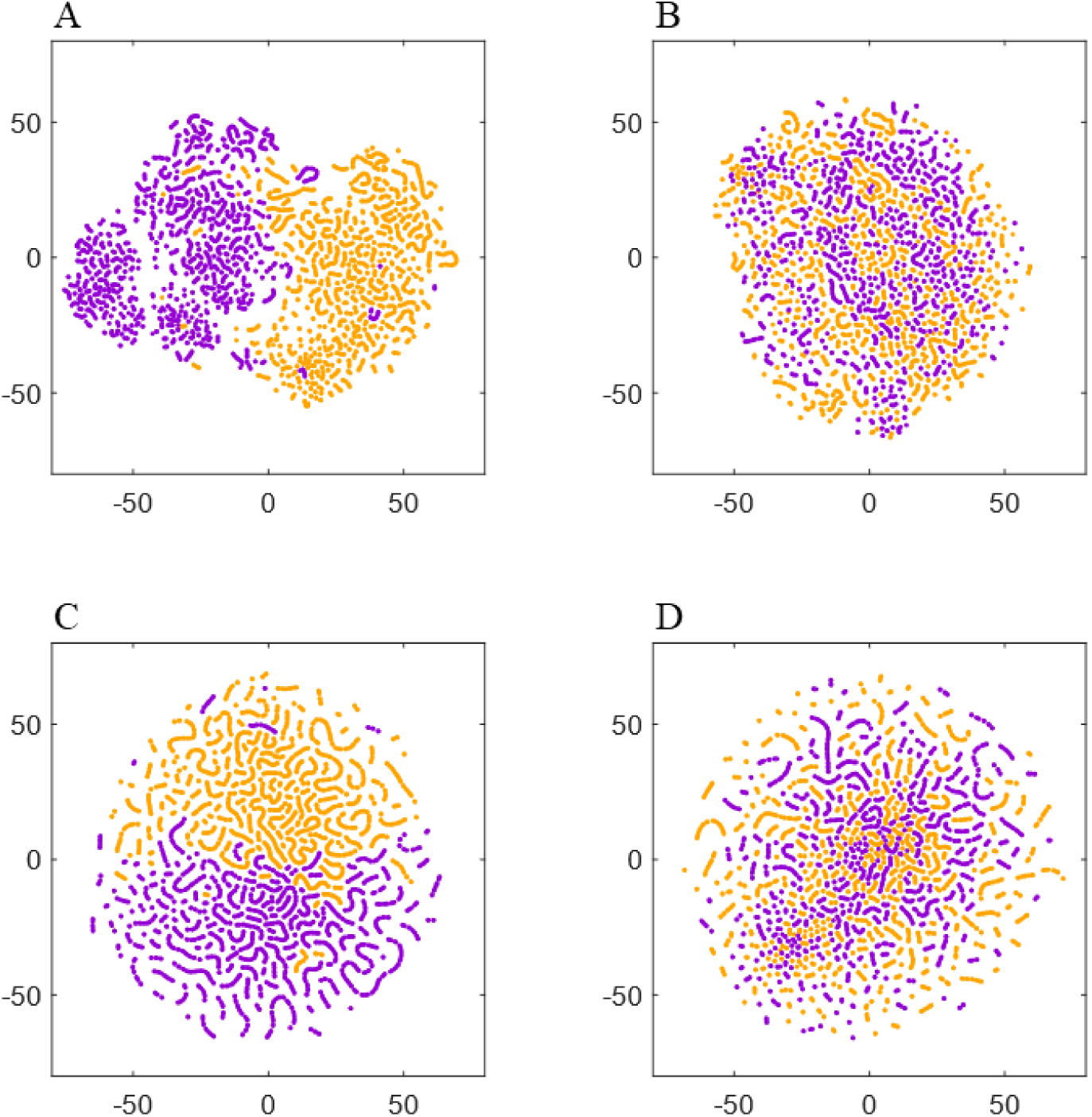
T-SNE plots for the feature sets of the participant with the best (A and C) and the worst (B and D) classification accuracy. Samples of the classes ‘Tenderness’ and ‘Anguish’ are plotted in orange and violet, respectively. The upper two plots (A and B) depict the datasets of the time-frequency analysis, while the lower two (C and D) display the amplitude time-series correlation features of the two participants.

Lastly, we extracted the twenty highest-ranked features for each subject and for each feature array to evaluate feature importance across participants. The main features of the time-frequency analysis are presented in Figure 6. Notably, the FAA coefficients did not occur in this selection. The extracted features showed the importance of the occipital channels O1, O2, and OZ, as well as temporal channels of both hemispheres. In the frontal site, we noticed a strong prominence of features in the left hemisphere compared to the corresponding right with a ratio of approximately 3:2 (99 to 62 features), when evaluating channels FP1/FP2, AF3/AF4, AF7/AF8, F1/F2, F3/F4, F5/F6, and F7/F8. We can report a particularly strong presence of channel AF7 in comparison to the complementary right-hemispheric channel AF8 as well as a strong dominance of gamma and high-beta bands in the highest-ranked features.

**Figure 6:**
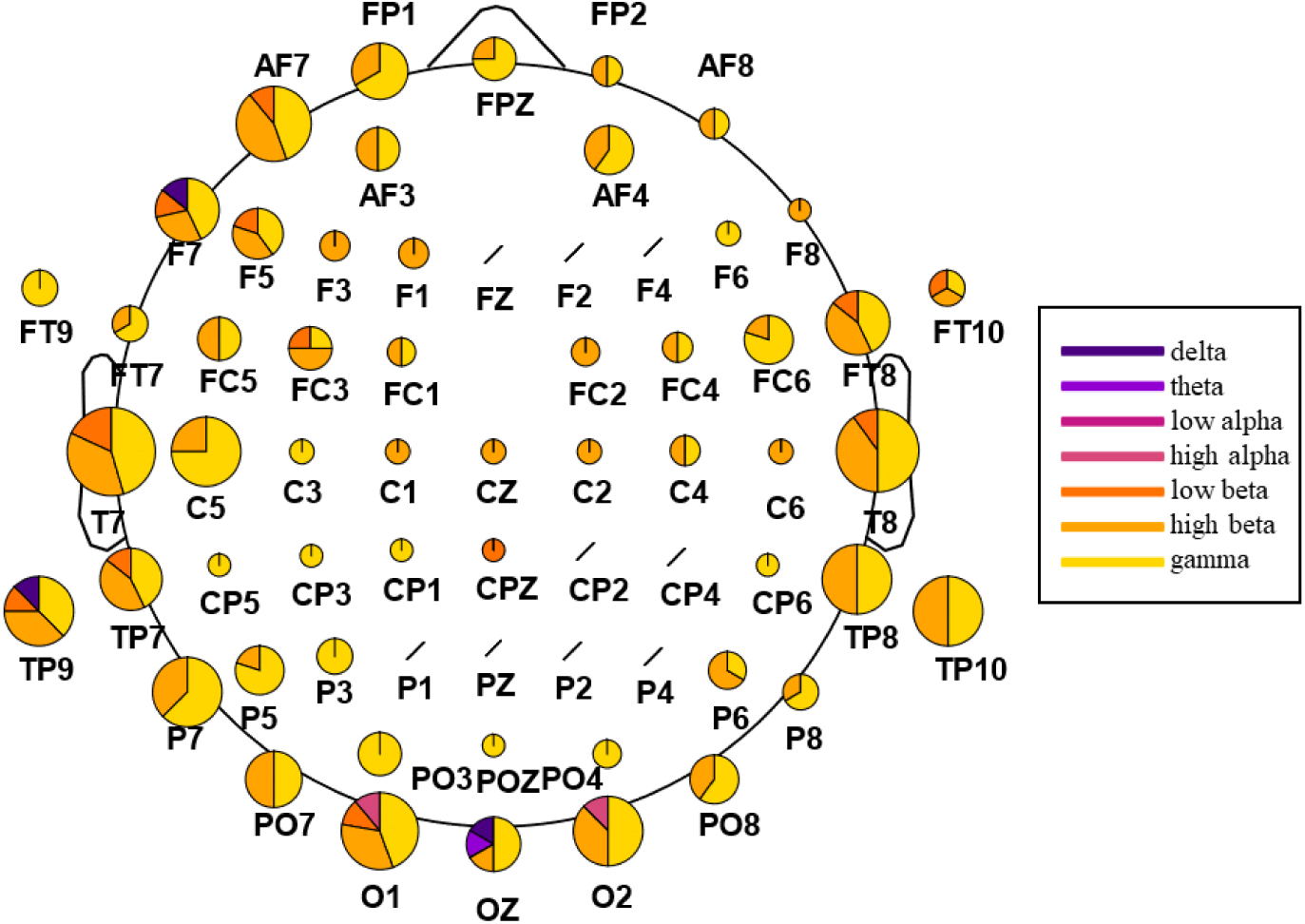
Extraction of the highest-ranked features for classification from the time-frequency analysis. The analysis was carried out across participants, with 220 features in total (20 per participant). The size of the channel-specific pie plot reflects how often the respective channel occurred in the main features. The distribution within the pie plots indicates the relative amount of features from each frequency band, ranging from delta (dark violet) to gamma (yellow). Channels marked with a black slash did not occur in the main features.

On the other side, the highest-ranked features extracted from the amplitude time-series correlational analysis appear to be more equally distributed across the seven analyzed frequency bands. In particular, features from theta and high-alpha ranges showed asymmetrical distribution in correlation, with a prominence in the left hemisphere for the former and in the right one for the latter. Significant connections in low-alpha and beta, especially high-beta, appear to be more prominent in the posterior part compared to the frontal one. The results also highlight the importance of temporoparietal and temporo-temporal correlations in the beta and gamma bands (Fig.7 E/F/G). In contrast, correlations in lower frequencies appear to be dominant in the frontal and occipital sites (Fig. 7 A/B/C/D). Taking together all frequency bands, we observe that correlations in the left frontal site occur with more prevalence across participants compared to connection in the right frontal hemisphere, with a ratio of approximately 2:1 (44 to 20 features) when comparing channels FP1/FP2, AF3/AF4, AF7/AF8, F1/F2, F3/F4, F5/F6, and F7/F8. Once more, we can report a high involvement of the left-hemispheric channel AF7 in comparison to the complementary right-hemispheric channel AF8, in particular in the high alpha band (Fig. 7D).

**Figure 7:**
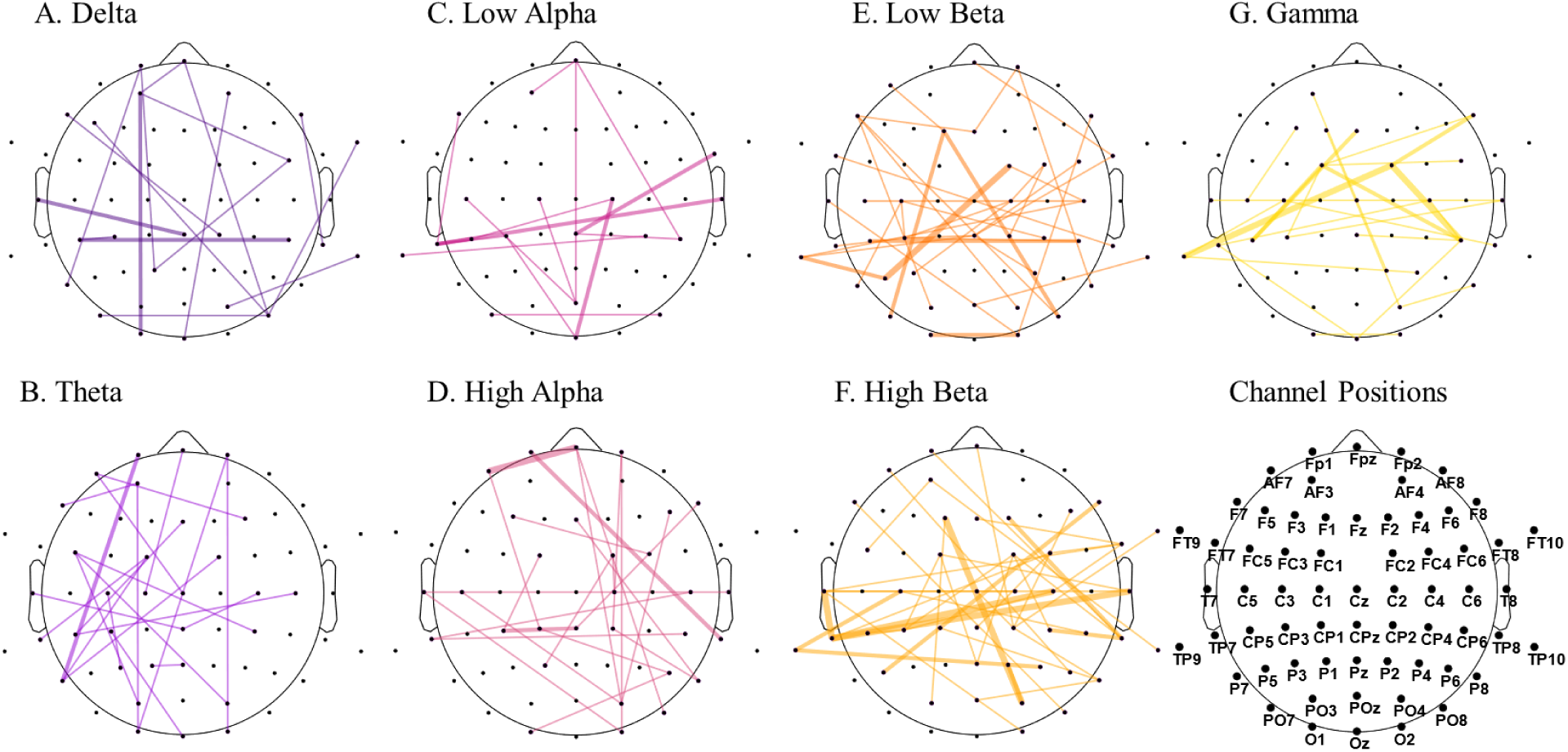
Features with the highest significance in distinguishing the emotional states from the amplitude time-series correlation analysis. The analysis was carried out across participants, with 220 features in total (20 per participant). The graph plots are separated into the seven extracted frequency bands, from delta (dark violet) to gamma (yellow). The width of the connection reflects how often the correlation occurs in the main features across participants, with the thicker line representing higher occurrence across subjects.

## Discussion

Our proof-of-concept study examined the feasibility of using EEG and machine-learning tools to distinguish complex emotional states successfully.

The results of the within-subject classification showed that it is possible to discriminate the electrophysiological correlates of two discrete, higher-order emotions, such as anguish and tenderness. We believe that this achievement is partly due to the way we evoked these emotions in the experimental setting. Firstly, the use of a multimodal realistic virtual environment may have eased and encouraged the participants’ involvement in the experiment by providing an engaging setup, as proven by previous studies (Kim et al., 2014; Cho et al., 2002; Lécuyer et al., 2008; Kovacevic et al.,2015). Secondly, the accompaniment of musical excerpts may have facilitated emotion-elicitation, given that music is a powerful tool for inducing strong emotional experience (Blood et al., 1999; Brown et al., 2004; Trost et al., 2012; Koelsch, 2014). Nevertheless, the usage of the same musical track may have influenced participants’ emotional elicitation differently due to variances in individual taste in music. For future work on emotion elicitation, choosing personalized musical tracks could improve individuals’ induction of emotional states. Lastly, the self-induction of emotional states through personalized mantras may have influenced the good outcome of discrete emotion classification, since we combined external elicitation with an internal locus of emotional induction. However, this may also have played a role in the substantial inter-subject variability in classification accuracy, given that self-reported ratings showed high variance in participants’ responses to the mantras’ usefulness.

Indeed, using spectral and FAA features we report accuracies ranging from 66.3% up to 95.6% when using random cross-validation, and 59.2% up to 92.9% when testing the model on unseen data of whole trials. Similarly, features extracted from the amplitude time-series correlation analysis showed a comparable model performance when tested on unknown data, with the lowest accuracy being 62.4% and the highest 92.4%, The nonergodicity in human subjects studies can explain the high interindividual variability we found related to these results (Fisher et al.,2018). Several studies have observed individual differences in emotional processing (Canli, 2004; Aftanas et al., 2006; Molenaar & Campbell,2009; Kuppens et al., 2009), stressing the importance of analyzing data at the individual level.

Notwithstanding, we assume that different amounts of noise in the dataset may have contributed to the high inter-individual variability of classification results. As stated in the methods section, we decided not to apply any artifact removal or epoch rejection going beyond ICA to remove eyeblink and muscular artifacts. We wanted to test the performance of the classifier as close as possible to the practical NFB scenario, in which epochs cannot be rejected either for training or for testing the model. We prioritized the utility of our model in real-world application and aimed for a robust classification, despite the inclusion of noisy window-samples and a conservative validation with samples from unseen trials.

Noticeably, the between-subjects variance in SVM model performance is significantly reduced when using correlational features and random cross-validation (91.6%-99.2%). We assume that it is because correlational features are likely to be stable across several adjacent 1-second windows, which suggests an underlying temporal structure of connectivity dynamics and a dependency in the sample set. The central assumption of random cross-validation, however, is that training and validation data are independent, or the estimated model accuracy will be too optimistic, and the model selected will be too complex (Roberts et al., 2017). In fact, when validating the model with entirely unknown trials, we observed a drop in accuracy of 22.3% on average for the classification with amplitude time-series analysis.

Supporting what stated above, the t-SNE plots in Figure 5 give insight into this temporal structure by grouping temporally adjacent samples closer together. This local pattern dominates over the global data structure for participants with low classification accuracies (Fig. 5D), suggesting that this phenomenon depends on temporal properties of connectivity rather than changes in neural correlation directly related to the emotional states. Nevertheless, our mean classification results of 73% for samples from trials unknown to the classifier (Table 1) and a precise formation of two global clusters for participants with high classification accuracy (Fig. 5A and C), confirm that both emotions show distinct electrophysiological correlates beyond these temporally changing patterns in neural activity.

Although of high importance for the prevention of overly positive classification results, strategic validation methods, such as trial-specific cross-validation, are sparsely used in fMRI and EEG time-series classification with sliding windows. In most studies, no alternative validation method is applied in addition to random cross-validation.

Surprisingly, the results of the feature selection algorithm when using the first set of features extracted from the time-frequency analysis did not show the importance of FAA features nor the midline spectral power as in previous classification studies involving tenderness (Zhao et al.,2018). However, as shown in Figure 5, the left frontal electrodes, especially channel AF7, appeared to have a substantial weight across participants on the model performance in discriminating between the two emotions. The relevance of the electrode AF7 is in line with previous studies using MUSE EEG headband comprising four channels (TP9, AF7, AF8, TP10) for emotion classification purposes, highlighting the importance of channel AF7 for accurately distinguishing between mental states (Bird et al., 2019; Raheel et al., 2019). Interesting to notice is that channels from the right frontal hemisphere did not appear to have such an impact across participants as much as their left-sided counterparts.

Regarding the contribution of the different frequency bands, high-beta and gamma bands resulted in having the highest impact on discrimination between emotional states across subjects. This result is also consistent with the literature providing evidence of the importance of these bands for distinguishing different emotional states (Müller et al., 1999; Keil et al., 2001; Glauser & Scherer 2008; Li & Lu, 2009; Daly et al., 2014).

However, it has to be noticed that we cannot extract any biological marker from this analysis since this feature-ranking algorithm does not allow us to infer which of the two classes showed increased or decreased spectral power or amplitude-related coherence patterns.

This study presents some pitfalls, most importantly, the small sample size and the non-randomization of trials across participants. Another limitation of our study, as already stressed above, is the high intersubject variability we found in the classification of the two affective states. Since emotions both at experiential and neurophysiological levels cannot be reduced to constant patterns across individuals (Barret et al., 2007; Kuppens et al., 2009), following studies should involve a bigger sample size. This will help shed light both on the individual differences and the common characteristics of emotional dynamics. Future directions would be to test this experimental protocol and SVM model on the training of such emotions through EEG-based neurofeedback, which can represent another tool for exploring these affective states. Besides its clinical application, NFB can be regarded as a perturbative approach in which the activation patterns used to give the feedback act as independent variables, thus enabling causal inferences to be made between brain activity and behavior (Sitaram et al., 2017).

In conclusion, the main contributions of this work are as follows:

- We provided evidence that complex emotions, such as anguish and tenderness, show distinct electrophysiological correlates.
- We demonstrated that, despite poor spatial resolution and the inability to infer activity in subcortical structures, EEG is a suitable tool for classifying discrete emotions, and that the proposed pipeline may be applicable for neurofeedback applications that require real-time data processing.
- We reported an asymmetrically high significance of both spectral and correlational features in the left frontal hemisphere compared to the right, particularly in channel AF7 compared to channel AF8.
- We discussed the importance of combining random cross-validation with strategic cross-validation methods when classifying time-series data.

## Acknowledgements

This study was supported by internal grants from the D’Or Institute for Research and Education (IDOR), and grants by the National Institute for Translational Neuroscience (INNT/Brazil) and the Fundaçao de Amparo à Pesquisa do Estado do Rio de Janeiro (FAPERJ; E-26/ 202.962/2015).

